# Attention to a threat-related feature does not interfere with concurrent attentive feature selection

**DOI:** 10.1101/356402

**Authors:** Maeve R. Boylan, Mia N. Kelly, Nina N. Thigpen, Andreas Keil

## Abstract

Visual features that are associated with a task and those that predict noxious events both prompt selectively heightened visuocortical responses. Conflicting views exist regarding how the competition between a task-related and a threat-related feature is resolved when they co-occur in time and space. Utilizing aversive differential Pavlovian conditioning, we investigated the visuocortical representation of two simultaneously presented, fully overlapping visual stimuli. Stimuli were isoluminant red and green random dot kinematograms (RDKs) which flickered at two tagging frequencies (8.57 Hz, 12 Hz) to elicit distinguishable steady-state visual evoked potentials (ssVEPs). Occasional coherent motion events prompted a motor response or predicted a noxious noise. These events occurred either in the green (task cue), the red (threat cue), or in both RDKs simultaneously. In an initial habituation phase, participants responded to coherent motion of the green RDK with a key press, but no loud noise was presented at any time. Here, selective amplification was seen for the task-relevant (green) RDK, but interference was observed when both RDKs simultaneously showed coherent motion. Upon pairing the threat cue with the noxious noise in the subsequent acquisition phase, the threat cue-evoked ssVEP (red RDK) was also amplified, but this amplification did not interact with amplification of the task cue, and did not alter the behavioral or visuocortical interference effect seen during simultaneous coherent motion. Results demonstrate that although competing feature conjunctions result in interference in visual cortex, the acquisition of a bias towards an individual threat-related feature does not result in additional cost effects.

## Significance statement

Selectively perceiving and adaptively responding to cues associated with danger are fundamental functions of the vertebrate brain. In humans, their disruption or dysregulation is at the core of many psychiatric diagnoses, including fear, anxiety, post-traumatic syndromes, and mood disorders. The present study examined the competitive interactions between the prioritization of threat cues and a concurrent cognitive task, to characterize how the human attention system manages limited resources in the presence of threat. Results showed that the selection of an individual feature signaling imminent threat is not at the cost of concurrent attention performance, even when threat and task stimuli overlapped in space. Findings support recent models of emotion/attention interactions that emphasize flexible, feature-based allocation of resources to biologically relevant stimuli.

## Introduction

The visual system receives dense sensory information, continually exceeding the limited capacity of visual cognition. In response to this challenge, the human brain has evolved mechanisms for selecting and prioritizing behaviorally relevant stimuli over other stimuli in the surrounding environment. Prioritization of task-relevant stimulus representations in the visual cortex has been documented extensively in research on selective attention (Hillyard et al., 1973; Reynolds and Heeger, 2009). A growing body of research has also demonstrated that stimuli associated with threat or danger prompt selective visuocortical amplification. For example, in aversive conditioning, visual cues paired with noxious outcomes elicit heightened visuocortical responses (Miskovic and Keil, 2012). Outside the laboratory however, observers confront visual environments in which task-relevant and threatening stimuli compete for limited capacity, prompting studies of emotional distraction and task interference (Mather and Sutherland, 2011; Iordan and Dolcos, 2015). This body of work has converged to show that naturalistic task-irrelevant distractors high in emotional intensity, such as mutilation scenes, interfere with performance at a concurrent task (Bradley et al., 2012) and diminish the visuocortical response elicited by the concurrent task stimuli (Müller et al., 2008). By contrast, studies in which individual threat-conditioned features such as orientation, color, or visual motion direction cooccur with spatially separated task cues have reported lack of interference effects (Miskovic and Keil, 2013, 2014). This is consistent with observations during feature-based selective attention, in which interference between competing features is restricted to the locus of spatial attention (Forschack et al., 2016).

The question arises regarding how threat-related features compete with task-relevant features for limited capacity. Specifically, do interference effects occur because of spatial competition alone, or is interference contingent on threat cues and task cues sharing non-spatial features, such as color, orientation, or motion direction? Addressing these questions has clinical implications, given increasing efforts to use measures of selective attention to threat for diagnostic assessment and intervention in a range of psychiatric disorders (e.g., Amir et al., 2010). Methodological barriers have hampered addressing this problem however: Most neurophysiological measurements cannot separately quantify the neural response to two simultaneously presented, overlapping stimuli. Steady-state visual evoked potential (ssVEP) frequency-tagging is a technique which yields direct, separate measurements of the visuocortical response evoked by individual items embedded in a complex array. The ssVEP is evoked when a visual stimulus is periodically modulated in terms of luminance or contrast (Norcia et al., 2015; Wieser et al., 2016). It can be extracted from scalp EEG, using spectral analysis, at the same fundamental frequency as the driving stimulus, often including higher harmonics. Critical for the present study, this technique separately quantifies the visuocortical response to concurrently presented and spatially fully overlapping stimuli.

Using this technique, we combined a feature-based attention task with an aversive conditioning protocol, using two fully overlapping RDKs differing in color, one being task-relevant, one threat-relevant: Occasional feature conjunctions of color and coherent motion in a given kinematogram represented a task cue or a threat cue, prompting a key press or predicting the onset of an aversive loud noise, respectively. Trials could contain task cues, threat cues, or both simultaneously. We quantified the frequency-tagged ssVEP evoked by each RDK during random and coherent motion, before and after conditioning, enabling us to test alternative hypotheses regarding the nature of threat-based selective attention. Hypotheses emphasizing spatial competition effects of threat (e.g., Holmes et al., 2003; Bishop, 2008) predict that selection of a threat cue should be at the cost of concurrent stimuli at the same location, resulting in a competitive trade-off between threat-evoked and task-evoked visual responses, during random as well as during coherent motion. Alternative perspectives have emphasized feature-specific rather than space-based amplification of threat representations (Pessoa and Adolphs, 2010; Mather, 2007). These notions predict that competitive interactions should be restricted to situations where threat and task features are similar (McTeague et al., 2015) or when conjunctions of more than one feature are present (Notebaert et al., 2011).

## Materials and Methods

### Participants

Seventeen undergraduate students (12 female, 3 left-handed; mean age = 19; SD = 1.7) with normal or corrected-to-normal vision and no personal or family history of photic epilepsy participated in this study. Participants received credits in a general psychology course for their participation. One participant was excluded from data analysis due to excessive EEG artifacts (more than 50% of trials were rejected). Experimental procedures were approved by the institutional review board at the University of Florida and were in accordance with the Declaration of Helsinki.

### Experimental Design and Stimuli

Visual stimuli consisted of one red and one green, isoluminant (see procedure), random dot kinematogram (RDK) presented simultaneously against a black background. A centrally located white fixation point was present throughout all trials and during the intertrial interval. Both RDKs spanned the same visual angle of 6.9° and consisted of 120 continuously moving dots (0.5° each), generated and presented using Matlab code and PsychToolbox functions (Brainard, 1997). Each dot within the RDK was displaced 0.2° in a random direction every 116 ms. During each trial, all dots of the RDK could systematically move in a coherent motion in one of eight cardinal directions (0°, 45°, 90°, 135°, 180°, 225°, 270°, 315°). Stimuli were presented centrally on a 23-inch Samsung SyncMaster SA950 LED monitor with a refresh rate of 120 frames per second, placed at a viewing distance of 150 cm.

An aversive differential Pavlovian conditioning paradigm with three phases (habituation, acquisition, extinction) and a total of 180 trials (60 trials per phase) was implemented. Each trial began with the simultaneous presentation of the overlapping red and green RDKs against a black background, and both RDKs remained on the screen for 8.17 seconds. Each trial represented one of three experimental conditions, which were determined by occasional coherent motion events (one event per trial) among otherwise randomly moving dots. Coherent motion events occurred randomly (drawn from a rectangular distribution) between 1750 ms and 6413 ms after RDK onset and lasted for 1749 ms. Thus, the first 1749 ms of each trial never contained coherent motion events. The three conditions were defined by which RDK displayed a coherent motion event in a given trial: Coherent motion of the green RDK represented the *target* condition, during which participants were instructed to respond with a button press. Coherent motion of the red RDK represented a conditioned *threat cue*, paired with a loud noise during the acquisition phase (see below). *Concurrent coherent motion* of both RDKs represented the third condition. The three conditions and the sequence of events for any given trial are depicted in Figure 1. Each condition was presented randomly 20 times during each phase of the experiment. Participants were instructed to detect the coherent motion event in the green RDK in the *target* and *concurrent* conditions, and to respond as quickly and accurately as possible by button press while ignoring coherent motion in the red RDK.

**Figure 1.**
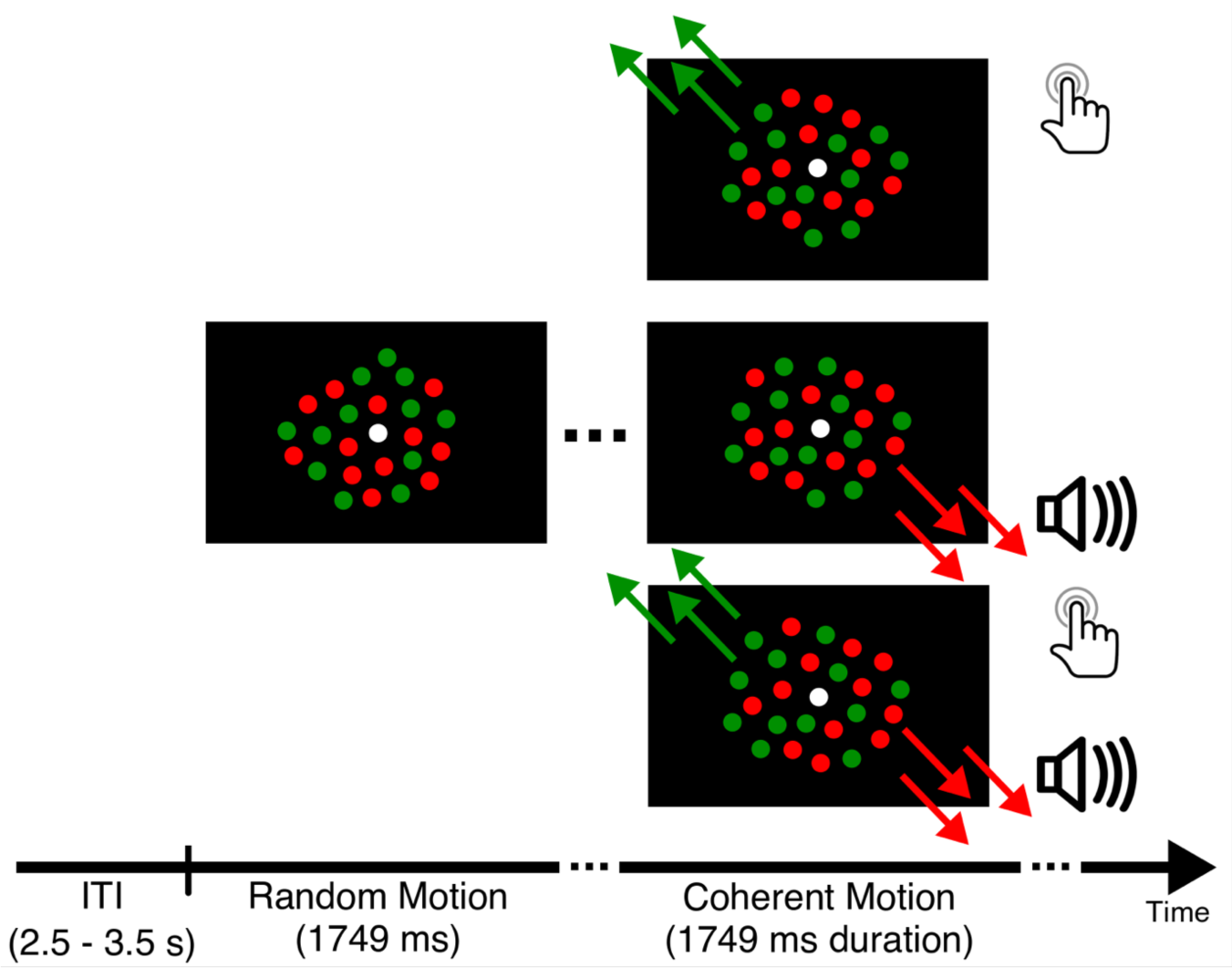
Trial sequence. For any given trial, during the first 1749 ms all dots presented on the screen moved in a random motion. Here, the only relevant feature was color, as participants were told to attend the task-relevant green dots. Following this guaranteed segment of random motion, all of the dots in one or both RDKs could move coherently in one direction. The onset of coherent motion could occur between 1750 ms and 6413 ms post-stimuli onset. Similar to the random motion segment, the coherent motion segment had a duration of 1749 ms. Regardless of experimental phase, when the task-relevant green dots exhibited coherent motion *(target* and *concurrent* conditions), participants responded via button press. During acquisition, the coherent motion of the red dots *(threat* and *concurrent* conditions) was paired with an aversive loud noise (the unconditioned stimulus). In total, the visual stimuli were presented for 8170 ms. The intertrial interval was 2.5-3.5 seconds, during which the participants viewed a black screen and a centrally located, white fixation point.

During the acquisition phase, coherent motion of the red RDK served as the conditioned stimulus (CS) and predicted the occurrence of the unconditioned stimulus (US), which was 500 ms of a 92 dB (SPL) white noise burst, generated in Matlab, played over speakers placed directly behind the participant. Specifically, coherent motion events in the red RDK, which occurred in the *threat* and *concurrent* conditions, were paired with the US for the final 500 ms of coherent motion, upon which the coherent motion event and the loud noise co-terminated. Conditions were randomized and fully balanced throughout all trials. Intertrial intervals randomly varied between 2.5 and 3.5 seconds (drawn from a rectangular distribution).

In addition to continuous motion, each dot of the RDKs flickered throughout the trial to evoke ssVEPs. For the first half of participants, the task-relevant green RDK flickered at 8.57 Hz while the red RDK flickered at 12 Hz. The tagging condition was counterbalanced for the second half of participants in order to control for confounds of flicker frequency (i.e. green flickered at 12 Hz and red flickered at 8.57 Hz). This resulted in two distinguishable ssVEP responses, one for the target stimulus and one for the threat stimulus.

### Procedure

After providing written consent, the isoluminance level of the red dots relative to the green dots was determined using flicker photometry. Using monochromatic circles (1° of visual angle) embedded in a gray (first step) or monochromatic (second step) field, each observer first adjusted the intensity of the red channel of the LED until no flicker was perceived between alternating red and gray backgrounds, and then adjusted the red against a monochromatic green background. Each observer’s final red value was used for the remainder of the experiment.

Participants were given instructions to focus on the green dots, while ignoring the red dots, and to click the mouse when the green dots moved together in a coherent motion. Participants were made aware that occasionally loud noises would occur in the experiment, but no additional information was given regarding contingencies. Participants were then fitted with the appropriate EEG sensor net and seated in a dimly lit and sound attenuated Faraday chamber. A chin rest was used to ensure the participant remained 150 cm away from the monitor for the duration of the study. In addition to oral instructions, participants viewed on-screen instructions before each experimental phase. Prior to and following the acquisition phase, each participant gave affective ratings on a scale of 1-9, of the red and green dots using the Self-Assessment Manikin (SAM; Lang, 1980). At the end of the experimental session, all participants were debriefed.

### Data Acquisition and Reduction

EEG was continuously recorded with a 257-channel sensor net at a sampling rate of 500 Hz, using an Electrical Geodesic amplifier with an input impedance of 200 MΩ. Impedances were kept below 60 kΩ. During recording, online filtering occurred by means of a 0.1 Hz high-pass and 80 Hz (3-dB points, respectively) low-pass elliptical filters. After recording, all data were converted to an average reference and further filtering occurred offline.

All channels were preprocessed by means of a 3 Hz high-pass filter (defined as the 1-dB point, Butterworth, 2^nd^ order filter) and a 30 Hz (3-dB point) low-pass filter (Butterworth, 14th order filter). Epochs were segmented 1 second pre- and 8 seconds post-stimuli (RDKs) onset. Artifacts were rejected based on statistical parameters of these epochs (absolute value, standard deviation, maximum of differences) across time points and for each channel, as proposed by Junghofer et al (2000). Across subjects for the *threat* condition, an average of 16.06 (SD=3.49), 14.31 (SD=3.75), and 14.13 (SD=3.67) trials were kept after artifact rejection for the habituation, acquisition, and extinction phases, respectively. For the *target* condition, an average of 15.00 (SD=4.56), 15.31 (SD=2.70), and 12.81 (SD=2.99), and for the *concurrent* condition, an average of 16.63 (SD=2.78), 14.81 (SD=3.87), and 12.56 (SD=3.29) trials were kept after artifact rejection for the habituation, acquisition, and extinction phases, respectively.

#### Steady state visual evoked potentials (ssVEPs)

Artifact-free epochs were averaged for each of the three conditions in the three phases. For each trial, two segments were analyzed: (1) random motion and (2) coherent motion. During each trial, following stimuli onset, there were 1750 ms of random dot motion prior to any possible onset of coherent motion. These data segments were averaged to result in the “random motion” ssVEP, which is sensitive to attentive bias towards one of the RDKs that does not depend on the occurrence of target events. In a subsequent step, the coherent motion segments (also of 1750 ms length) occurring at different latencies in each trial were aligned relative to the beginning of the coherent motion and averaged, yielding the “coherent motion” ssVEP. Using inhouse Matlab code, these ssVEP signals were subjected to a Fourier transform to quantify the power at the driving frequencies (8.57 Hz and 12 Hz) of each of the two RDKs. This analytic step involved tapering the data with a 40-ms cosine-square window, followed by discrete Fourier transform across all 1750 ms, and normalizing the resulting complex spectrum by the number of points. The resulting power spectrum had a resolution of 0.57 Hz. To enable comparison between tagging frequencies and to eliminate the effect of nonspecific level differences between participants and conditions, we computed the signal-to-noise ratio (SNR) for each tagging frequency: We divided the power at each tagging frequency by the mean of the spectral power at 6 adjacent frequency bins in each spectrum, leaving out the two immediate neighbors.

### Statistical Analysis

#### Behavioral Data

Hit rate, false alarm rate, and response time were considered. The hit rate was the percentage of correct responses, defined as responses following detection of the coherent motion of the task RDK in the *target* and *concurrent* conditions between 150 and 2000 ms post-onset. The false alarm rate was the percentage of incorrect button presses when the task RDK did not exhibit coherent motion, and responses outside the “correct” window, defined above. Response time was the amount of time that elapsed between the onset of the coherent motion of the task RDK in the *target* and *concurrent* conditions and the button press for a correct response. The Self-Assessment Manikin (SAM) was used to collect affective ratings (hedonic valence) for the red and green dots, before and after the acquisition phase for each participant.

A repeated-measures ANOVA was conducted for the hit rate with phase (habituation, acquisition, extinction) and condition (target, concurrent) as factors. Likewise, a repeated-measures ANOVA was conducted for the response time with phase (habituation, acquisition, extinction) and condition (target, concurrent) as factors. Significant main effects or interactions were followed by post-hoc ANOVA and paired t-tests, as appropriate.

A repeated-measures ANOVA was conducted for the SAM ratings with conditioning (pre, post) and color (red, green) as factors. Significant main effects or interactions were followed by post-hoc ANOVA and paired t-tests, as appropriate.

Throughout this report, frequentist statistics were supplemented by Bayesian repeated-measures ANOVAs, where appropriate, to quantify the evidence for the different hypotheses as well as for null-findings. In each Bayesian analysis, the prior model probabilities were uniformly distributed such that each model was given equal weight (i.e. in a Bayesian repeated-measures ANOVA with five candidate models, each held a prior probability of 0.2). Bayesian repeated-measures ANOVAs were conducted by utilizing JASP software (v 0.8.6; JASP Team, 2018). Interpretation of Bayes Factors follows language guidelines developed by Jeffreys (1961).

#### ssVEPs

Replicating a large body of research (reviewed in Wieser et al., 2016), maximum amplitudes were observed at occipital electrodes. Figure 2 shows the grand mean ssVEP time series obtained at a sensor corresponding to location Oz of the international 10-20 system, and the grand mean topographical distribution of the power at the driving frequency. For statistical analysis, SNR values were averaged over a cluster of occipital electrodes, including Oz and 21 of its nearest neighbors (EGI sensors: 116-119, 124-128, 136-140, 147-150, and 156-159, see Figure 2). To investigate the ssVEP amplitude changes when only color information was present (random motion) as opposed to when a feature conjunction of color and motion was present (coherent motion segment) the SNR values from the two segments entered the statistical analysis (below) separately.

**Figure 2.**
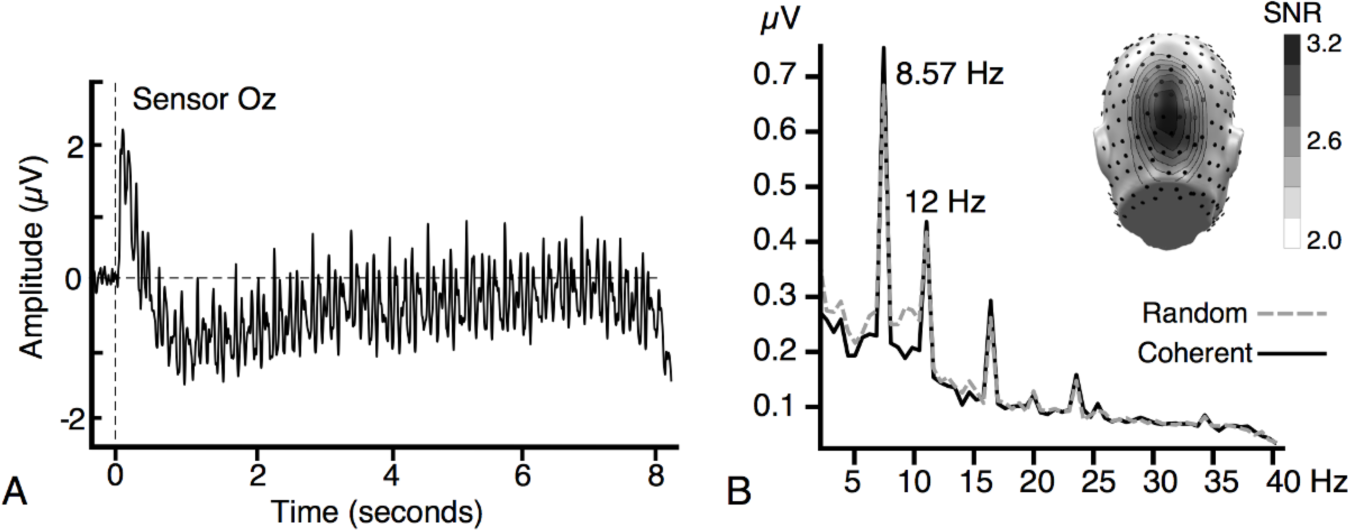
Time course and frequency spectrum. **(A)** Averaged time course for all trials and all participants extracted from sensor Oz. The vertical dashed line indicates stimuli onset. **(B)** Resulting frequency spectrum following a Fourier transform. Here, there are distinct peaks at the driving frequencies, 8.57 Hz and 12 Hz. Similar to previous ssVEP research, maximal amplitude power is found at occipital sensors (inset).

### Manipulation Checks

Before conducting targeted analyses of interaction between tagging frequencies, we conducted several manipulation checks to quantify the extent to which expected and established effects were present in this dataset. A repeated-measures ANOVA was conducted for only the *target* condition (coherent motion in the green RDK), with segment (random motion, coherent motion) and phase (habituation, acquisition, extinction) as factors, for the tagging frequency of the target stimulus. In this analysis, task-related enhancement of ssVEP amplitude would be reflected in a main effect of segment, indicating heightened ssVEP during target processing across all phases. To ensure that any such amplitude enhancement was specific to the target tagging frequency, the same repeated-measures ANOVA was conducted for the tagging frequency of the threat stimulus, where no amplitude enhancement during the target segment was expected. This comparison addresses potential concerns regarding the effects of transient brain events on ssVEP amplitude. If such transient events would contaminate the present ssVEP measures, then they are expected to affect both tagging frequencies, rather than one tagging frequency selectively.

In the same vein, we conducted a manipulation check on the red RDK, examining the extent to which the present data replicated the known selective amplification of the ssVEP evoked by conditioned threat cues. To investigate this effect, a repeated-measures ANOVA was conducted only for the *threat* condition (coherent motion in the red RDK) with segment (random motion, coherent motion) and phase (habituation, acquisition, extinction) as factors. Here, the expected amplification of the threat cue during acquisition would result in a selective amplification of the ssVEP evoked by the red RDK during acquisition. The specificity of any such effect to the threat cue would be demonstrated by the absence of this phase effect in the green RDK.

### Hypothesis Testing

To examine the central hypotheses of this study regarding the nature of trade-off and interference effects between the two RDKs, repeated-measures ANOVAs were conducted for random motion and for coherent motion segments separately:

For the random motion segment, a repeated-measures ANOVA was conducted with tagging frequency (threat stimulus, target stimulus) and phase (habituation, acquisition, extinction) as factors. Here, the experimental conditions in each phase were averaged together, because during random dot motion the trial condition (the type of coherent motion occurring afterwards) was irrelevant. A post-hoc ANOVA and paired t-tests were used to follow up significant interaction, where appropriate.

For the coherent motion segment, a repeated-measures ANOVA was conducted for the target stimulus tagging frequency with phase (habituation, acquisition, extinction) and condition (threat, target, concurrent) as factors. Here, an interaction between phase and condition would indicate an effect of fear acquisition on target stimulus processing, which would be consistent with competitive cost effects. Paired samples t-tests were used to follow up significant results, as appropriate. Additionally, Bayesian repeated-measures ANOVAs were conducted where appropriate. As an added control, the same ANOVA design was evaluated for the threat cue tagging frequency.

#### Competition Index

To directly quantify visuocortical competition between the two RDK stimuli, a competition index was calculated which represented the product of the amplitude changes from habituation to acquisition and extinction within each tagging frequency:

1. (Threat ssVEP_Acquisition_ − Threat ssVEP_Habituation_) * (Task ssVEP_Acquisition_ − Task ssVEP_Habituation_)
2. (Threat ssVEP_Extinction_ − Threat ssVEP_Habituation_) * (Task ssVEP_Extinction_ − Task ssVEP_Habituation_)

This index is negative if amplitude enhancement in the threat RDK is accompanied by amplitude reduction in the task RDK, or vice-versa (trade-off), but will be positive if both RDKs display amplitude reduction or amplitude enhancement from habituation to the subsequent acquisition or extinction phases. For statistical analysis, a repeated-measures ANOVA was conducted on the competition index with phase (difference between habituation and acquisition, difference between habituation and extinction) and condition (threat, target, concurrent) as factors for both the random motion segment and the coherent motion segment.

## Results

### Behavioral Data

Across participants, the overall average hit rate was 89.43% (SD = 19.22) and the average response time was 1662 ms (SD = 336). The average false alarm rates were 9%, 6%, and 3% for the habituation, acquisition, and extinction phases, respectively. Hit rate and response time, averaged across participants by condition are reported in Table 1.

**Table 1.**
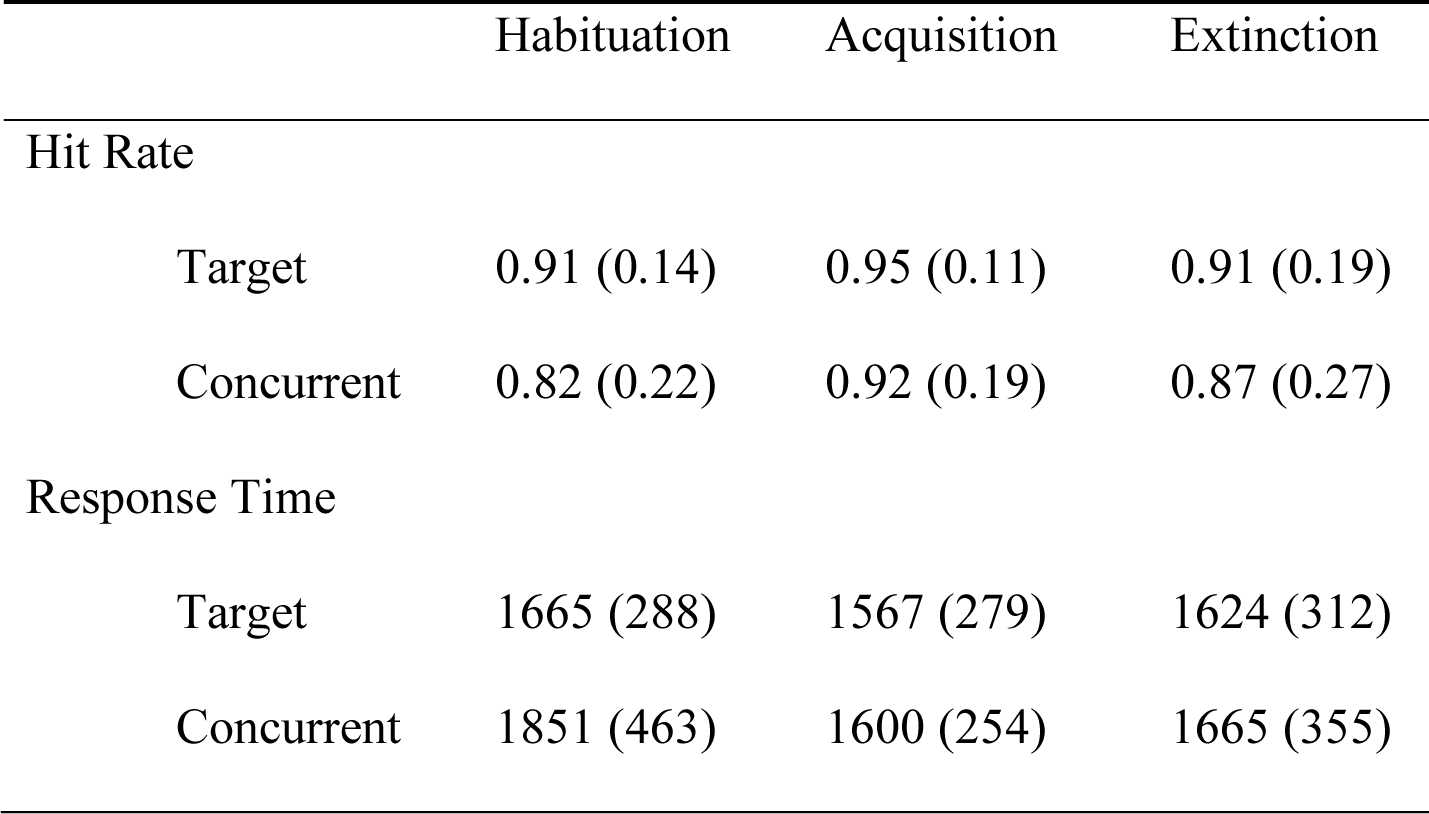
Means (SD) of hit rate and response time (in milliseconds) averaged across participants by condition and phase

The hit rate was not sensitive to any experimental manipulations, resulting in no significant main effects nor a significant interaction between phase and condition. By contrast, a significant interaction between phase and condition emerged for the response time (F(2,30) = 4.660, p = .021; η^2^ = .237). This interaction was driven by the significantly longer response time for the *concurrent* condition in the habituation phase compared to the other conditions, across phases (t15 > −2.962, p < .018, see Figure 3). The Bayesian repeated-measures ANOVA on response time resulted in a BF10 > 2.8 for the main effect of condition model with all remaining models returning a BF10 > 104.6, providing anecdotal to decisive evidence, respectively, for the alternate hypothesis. The model which received the most support was the two main effects (phase + condition) model with a BF10 of 485.2.

**Figure 3.**
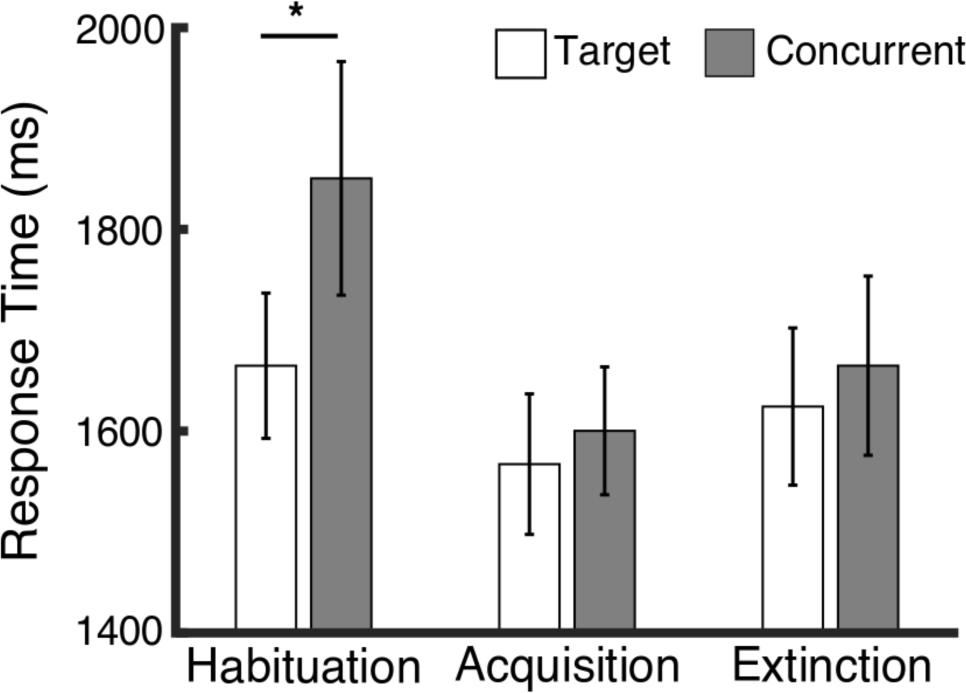
Response Time. Participants were slower in responding to the target cue during the *concurrent* condition in which both RDKs exhibited coherent motion. While this interference effect was pronounced during habituation it was not sustained throughout the acquisition or extinction phases of this experiment.

The SAM hedonic valence ratings were sensitive to experimental manipulation, resulting in a significant interaction which was driven by the red dots’ affective rating significantly increasing from pre-conditioning to post-conditioning (t_15_ = −6.603, p < .001) and a significantly higher SAM rating for the red dots than the green dots after conditioning (t_15_ = −5.883, p < .001). The average SAM rating before conditioning was 4.4 (SD = .66). Following conditioning the SAM ratings for the green dots did not significantly change (M = 4.3, SD = .54), while the SAM rating for the fear-conditioned red dots significantly increased (M = 6.0, SD = .78). The Bayesian ANOVA model which received the most support was the interaction model with main effects (conditioning + color + conditioning*color) with a BF10 > 1,000,000.

### Steady state visual evoked potentials (ssVEPs)

#### Manipulation Checks

Visuocortical responses to the task stimulus during the coherent motion of the target RDK (Figure 4A) increased from the random motion to the coherent motion segment across all phases of the experiment (F(1,15) = 5.759, p = .030; η^2^ = .277). The Bayesian repeated-measures ANOVA resulted in a BF_10_ = 10.4 for the segment main effect model, providing strong evidence for the hypothesis that target events increase the ssVEP amplitude evoked by the task stimulus. The visuocortical response to the threat stimulus during the same coherent motion was not affected by the target event, i.e., it did not show an increase between segments or across phases. Thus, the target event prompted a selective increase in ssVEP amplitude that was specific to the time period of the target event, and to the task stimulus.

The visuocortical response to the threat stimulus during the coherent motion of the threat cue (Figure 4B) demonstrated a main effect of phase (F(2,30) = 4.955, p = .032; η^2^ = .248) which was driven by the significant increase in ssVEP amplitude from the habituation phase to the acquisition phase (t15 = −2.692, p = .017) whereas the increase from habituation to extinction was not significant (t15 = −1.943, p = .071). The Bayesian repeated-measures ANOVA resulted in a BF 10 = 2.8 for the phase main effect model as well as a BF10 = 4.0 for the two main effects (segment + phase) model, providing substantial evidence for the hypothesis that conditioned threat cues result in selective visuocortical amplification. In a control analysis, the visuocortical response to the task stimulus during the coherent motion of the threat RDK did not show an increase either between segments or across phases.

**Figure 4.**
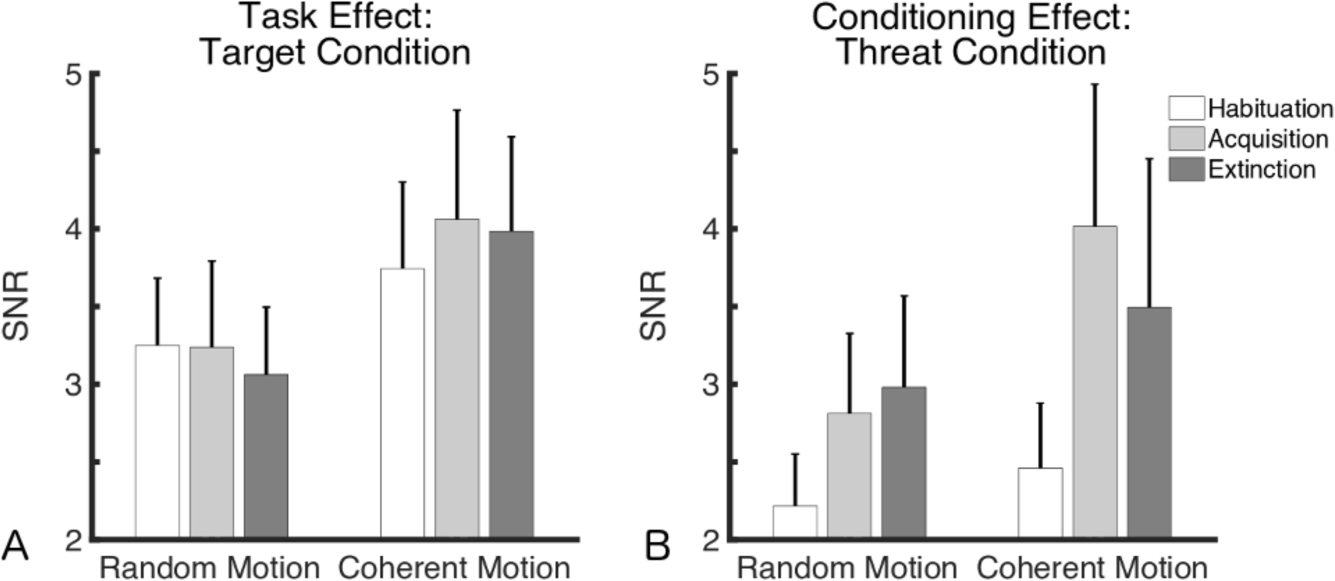
Manipulation Checks. **(A)** Visuocortical response for the target stimulus (i.e. the ssVEP for the task-relevant green dots) was amplified during the coherent motion segment, regardless of block, relative to the random motion segment. **(B)** Visuocortical response for the threat stimulus showed amplification following aversive conditioning. These findings replicate previous work in regards to selective amplification of both a task-relevant stimulus and a motivationally relevant, fear-conditioned stimulus in the visual brain.

#### Random Motion Segment

When the conditions within each phase were averaged for the segment of random motion, the ssVEP amplitudes during the first 1750 ms of each trial showed a significant interaction between tagging frequency and phase (F(2,30) = 4.587, p = .039; η^2^ = .234). A post-hoc ANOVA was conducted for both tagging frequencies and showed a main effect of phase for the threat tagging frequency (F(2,30) = 4.770, p = .028; η^2^ = .241). This interaction and main effect was driven by the significant increase in the threat stimulus ssVEP amplitude from habituation to acquisition (t15 = −2.498, p = .025, see Figure 5, top panel) and from habituation to extinction (t15 = −2.258, p = .039). The Bayesian repeated-measures ANOVA resulted in a BF10 = 3.5 for the tagging frequency main effect model, providing substantial evidence for the alternate hypothesis.

**Figure 5.**
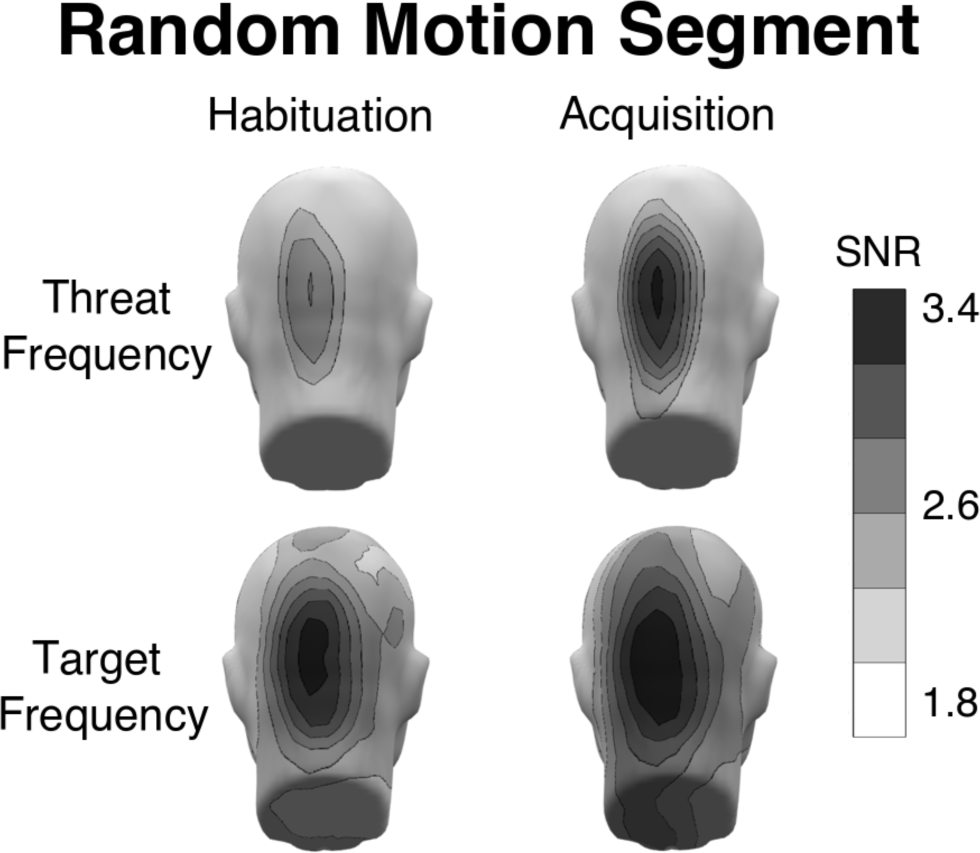
Random Motion Segment. During the random motion segment, the ssVEP for the threat stimulus showed significant amplification during acquisition, when coherent motion of the red RDK was paired with the unconditioned stimulus. The target stimulus did not prompt a similar amplification during acquisition but had relatively higher visuocortical activity throughout the experiment.

#### Coherent Motion Segment

The visuocortical response to the target stimulus was sensitive to the experimental conditions, (F(2,30) = 6.290, p = .008; η^2^ = .295, see Figure 6). Throughout the experiment, ssVEP amplitude was heightened during the coherent motion event during the *target* condition, compared to both the *threat* condition (t15 = −4.463, p = .000) and the *concurrent coherent motion* condition (t15 = 2.616, p = .019). The Bayesian repeated-measures ANOVA resulted in a BF_10_ = 2.7 for the main effect of condition model, providing anecdotal evidence for the hypothesis that coherent motion prompts selective amplification of the target response. Results of the ANOVA evaluating the threat stimulus tagging frequency converged with those of the manipulation checks, demonstrating heightened visuocortical activity for the conditioned threat stimulus during the acquisition phase.

**Figure 6.**
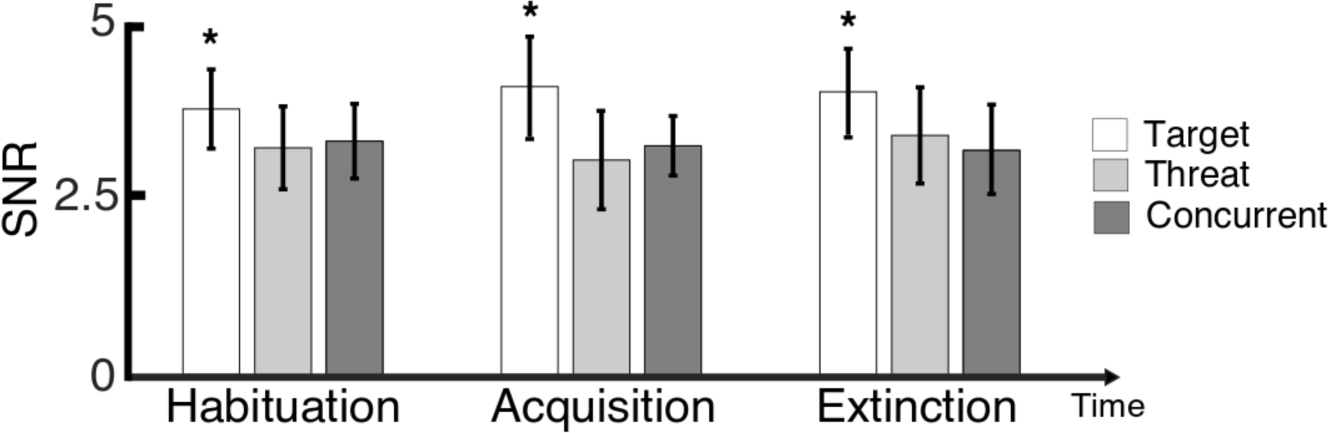
Coherent Motion Segment. The ssVEP for the target stimulus demonstrated a main effect of condition, in which the *target* condition had relatively higher amplitude for every phase of the experiment. Here, an interaction effect would indicate cost effects as the threat stimulus becomes relevant to the participant through fear conditioning. No interaction was found, suggesting that the amplification of the threat stimulus is not at the cost of the simultaneously presented target stimulus.

#### Competition Analysis

The repeated-measures ANOVA for the random motion segment and the coherent motion segment each resulted in no significant interaction between phase and condition, nor any main effects, suggesting that there is no visuocortical competition between the two RDKs throughout all phases of the experiment. The Bayesian repeated-measures ANOVAs for the competition index during coherent motion and random motion resulted in a BF01 > 3.0 and a BF01 > 4.6, respectively, for all models, providing substantial to very strong evidence for the *null* hypothesis. The two main effects with the interaction (phase + condition + phase*condition) models for both the coherent motion and random motion received the most support for the null hypothesis with BF01 of 31.8 and 77.2, respectively.

## Discussion

The present study asked if acquired biases towards a threat-related feature are at the cost of concurrent task-based feature selection at the same location in the visual field. We tested the alternative hypotheses that (1) threat cues interfere with concurrent stimuli at the same location irrespective of their feature composition and (2) that individual threat features are amplified without imparting cost effects on concurrent task-based feature selection, even when competing for representation at the same spatial locations. We addressed these hypotheses using the unique capability of ssVEP frequency tagging to quantify distinct neural signals from fully overlapping stimuli.

Combining a feature-based attention task with an aversive conditioning protocol, we replicated previous findings that a task-relevant feature prompts selectively heightened visuocortical responses (Muller et al., 2006). We also replicated previous work showing visuocortical response amplification following aversive conditioning (Moratti and Keil, 2009). This selective amplification effect was not accompanied by attenuation of the task-evoked ssVEP, in either a pre-cue random motion segment, or during the time segment in which coherent motion occurred. Targeted analyses using a sensitive competition index supported the conclusion that the visuocortical amplification of an individual threat feature is not at the cost of concurrent feature selection at the same location. By contrast, co-occurring feature conjunction events (color and coherent motion in both RDKs simultaneously) prompted interference, and this interference occurred independently of aversive conditioning, present already in the initial habituation stage. Visuocortical interference during simultaneous feature conjunction events effect was consistent with response time data, which also indicated interference when coherent motion events co-occurred in both RDKs during the habituation phase. Notably, this behavioral interference effect diminished across phases and was absent during acquisition and extinction, providing further evidence against the notion that fear conditioning of an individual salient feature heightens competition.

Together, these results are in support of conceptual models of threat-based selection that emphasize the flexible allocation of resources and the effective usage of the massively parallel machinery of the visual system in support of survival (Pessoa and Adolphs, 2010). The present study adds to such models by showing that an acquired attention bias towards a threat feature is constrained by the specific capacity limitations of the visual system, which differ for feature-based and space-based selection (Andersen et al., 2008).

Future work may further investigate the neurophysiological mechanisms that underlie the translation of broad defensive engagement of the organism into feature-specific gain modulation at the level of visual neurons. Previous research has emphasized the role of re-entrant feedback signals from limbic structures such as the amygdaloid body to visual cortical areas in amplifying visual threat cues (Anderson and Phelps, 2001; Amaral et al., 2003). Although these connections are well established in primate models, including their directional properties, evidence of their involvement in the selective visual processing of threat has been scarce. Emphasizing norepinephrinergic modulation of perception and attention, the arousal-biased competition model (Mather and Sutherland, 2011) states that motivational states, such as avoidance of threat, bias neural competition and that autonomic/emotional arousal will increase the competitive advantage for motivationally relevant stimuli while simultaneously decreasing the priority of motivationally irrelevant stimuli. The present study suggests that such competitive interactions are most pronounced in situations in which threat cues are defined by multiple features, again paralleling canonical findings in selective attention and visual search (Treisman and Kanwisher, 1998). In line with this notion, recent evidence from concurrent EEG-fMRI recordings during aversive Pavlovian conditioning suggests that the same cortical regions associated with spatial and feature-based attention may also be involved in the selection of threat cue representations (Petro et al., 2017). Thus, multimodal neuroimaging with the present experimental design may allow refining and testing of the hypothesis that fronto-parietal attention networks under the control of limbic circuits mediate the selective visuocortical facilitation of threat (Yin et al., 2018).

A longstanding debate in the field of emotion/attention interactions has discussed the extent to which threat cues are processed in an automatic, or parallel fashion, with no or little cost to concurrent neurocognitive processes (Ohman and Mineka, 2001; Pessoa and Adolphs, 2010; Pourtois et al., 2013). The present study suggests that for conditioned stimuli, such parallel processing is seen when a threat cue is completely distinct from the features that characterize the concurrent stimuli, i.e., when the threat cue is defined by a unique, salient feature. No prioritization however was observed when two concurrent feature conjunctions were present. This is consistent with the notion that the effect of color as a feature known to readily guide visual search (Wolfe and Horowitz, 2017) may be amplified by stimulus value, further facilitating attentive selection.

In conclusion, the present study used a novel experimental design, which allows controlling the physical stimulus properties of two fully overlapping stimuli, one of which was associated with a noxious outcome. Results strongly supported the hypothesis that selective attention to an individual, salient feature associated with threat is not at the cost of concurrent attentive processing in healthy adult observers. The practical implications of this finding are manifold: For diagnostic assessment of psychopathologies in the fear, anxiety, and depression spectrum, the presence of interference effects in some patients may represent a marker of dysfunctional attention biases (Bar-Haim et al., 2007). For intervention, the online feedback of ssVEP amplitudes is routinely accomplished in brain-computer-interfaces (Muller-Putz and Pfurtscheller, 2008), and may thus be used to alter maladaptive attentive selection of threat.

## Acknowledgements

This work was supported by the National Institutes of Health, Grant R01 MH097320 to Andreas Keil.

